# sPLA_2_ inhibitor-loaded nanoparticles mitigate spontaneous osteoarthritis in guinea pigs

**DOI:** 10.64898/2026.10.30.763253

**Authors:** Qiushi Liang, Qi He, Yiyang Xu, Yijun Dai, Courtney Nuss, Jeremy D. Eekhoff, Sita Awasthi, Jiahao Liang, Tingfang Sun, Yanhua Lan, Michael P. Duffy, Varad Bhangui, Andrew Tsourkas, Zhiliang Cheng, Ling Qin

## Abstract

Osteoarthritis (OA) is a prevalent degenerative joint disease lacking effective disease-modifying therapies. Secreted phospholipase A2 (sPLA_2_), a key upstream regulator of inflammatory lipid mediators, has been implicated in OA pathogenesis, but its therapeutic potential in age-related OA remains unclear. Here, we evaluated the role of sPLA_2_ and the efficacy of nanoparticle-mediated sPLA_2_ inhibition in a spontaneous OA model using female Dunkin-Hartley (DH) guinea pigs. We first characterized OA progression and observed age-dependent increases in cartilage degeneration, synovitis, oxidative stress, and sPLA_2_ expression and enzymatic activity. We then developed sPLA_2_ inhibitor (thioetheramide-PC)-loaded nanoparticles (sPLA_2_i-NPs) and demonstrated their superior ability, compared to free inhibitor, to prevent IL-1β-induced cartilage degeneration in guinea pig cartilage explants. In vivo, intra-articular administration of sPLA_2_i-NPs from 3 months of age significantly attenuated cartilage destruction, synovial inflammation, and osteophyte formation in 7- and 10-month-old guinea pigs. These structural improvements were accompanied by normalization of anabolic and catabolic gene expression and a marked reduction in synovial fluid sPLA_2_ activity. Importantly, longitudinal gait analysis revealed that sPLA_2_i-NP treatment preserved limb loading, propulsion, and locomotor performance, demonstrating functional recovery. No systemic toxicity was observed following long-term treatment. Collectively, our findings identify sPLA_2_ as a key driver of spontaneous OA and establish nanoparticle-mediated sPLA_2_ inhibition as a promising disease-modifying strategy for age-related OA.

## Introduction

Osteoarthritis (OA) is the most prevalent degenerative disorder of the musculoskeletal system, affecting hundreds of millions of people worldwide.(1) The etiology of OA is multi-factorial, including joint injury, obesity, aging, and genetic predisposition.(2) As the disease progresses, patients suffer from joint pain, stiffness, deformity, and functional impairment, leading to reduced quality of life and shortened life expectancy. Currently, there is no effective drug treatment blocking or reversing the progression of this disease and developing novel OA therapies is of high clinical importance.

OA is accompanied by chronic, unresolved low-grade inflammation. Analyses of human OA tissues and various animal OA models have implicated a broad array of inflammatory factors as key contributors to OA pathogenesis.(3) Despite this strong association, anti-inflammatory strategies have shown limited disease-modifying effects. Non-steroidal anti-inflammatory drugs (NSAIDs) provide symptomatic relief but do not attenuate the structural progression of OA.(4) Moreover, systemic blockade of conventional inflammatory cytokines effective in rheumatoid arthritis (RA), such as anti-TNF or anti-IL-1β therapies, has failed to demonstrate meaningful clinical benefit in OA.(5)

Over the past several years, we have explored the possibility of targeting another inflammatory factor for OA treatment.(6, 7) Secreted phospholipase A_2_ (sPLA_2_) is a family of small, secreted enzymes that cleave membrane phospholipids at the sn-2 position to release free fatty acids (often arachidonic acid) and lysophospholipids. These bioproducts serve as precursors for inflammatory mediators in many chronic inflammatory diseases.(8, 9) It was previously reported that human OA patients exhibit a high sPLA_2_ activity or expression level in the synovial membrane, synovial fluid, and articular cartilage.(10–13) By examining human and mouse OA cartilage samples, we have shown that the amount of sPLA_2_ drastically increases when OA initiates and remains at a high level throughout OA progression.(6) Its inhibitor, thioetheramide-PC, effectively reduces cartilage degeneration in mouse OA cartilage explants.(6) To develop an sPLA_2_-responsive and nanoparticle (NP)-based interventional platform for OA management, we engineered sPLA_2_i-loaded NPs (sPLA_2_i-NPs) by incorporating either thioetheramide-PC or varespladib into the phospholipid membrane of micelles. Subsequent mouse experiments demonstrated the efficacy of sPLA_2_i-NPs in mitigating OA progression.(6, 7)

While mouse models are commonly used in OA research, various interventions are required to induce OA. In our previous sPLA_2_i-NP study, we utilized a surgical approach (destabilization of medial meniscus, DMM) and a non-invasive approach (mechanical loading) to induce OA in mice.(6, 7) The resultant OA phenotypes are more similar to post-traumatic OA (PTOA) but not age-related OA. While aged mice (over 20-month-old) develop progressive articular cartilage thinning and proteoglycan loss, their OA phenotype is usually very mild.(14–16) Together with the high mortality of mice at this stage, aged mice are not an ideal model for nontraumatic, idiopathic OA, the most common form of human OA.

By contrast, Dunkin-Hartley (DH) guinea pigs have been well characterized as a superior naturally occurring OA model.(17) These animals have OA onset as early as 3 months of age in the absence of any experimental intervention. The phenotype closely resembles human spontaneous OA: cartilage degeneration occurs predominantly in the medial compartment of the knee;(18) inflammatory cytokines and matrix-degrading proteases are significantly upregulated;(19–21) early-stage disease is marked by synovial inflammation, chondrocyte apoptosis, proteoglycan depletion, and cartilage fibrillation,(22, 23) whereas advanced disease is characterized by extensive cartilage erosion, osteophyte formation, and subchondral bone sclerosis.(23–25) Notably, OA progression in this model is bilaterally symmetric, enabling paired comparisons between the two knees within the same animal.(18)

In this study, we first confirmed OA progression and the increase of sPLA_2_ inflammatory signaling in the knee joints of DH guinea pigs. We then synthesized sPLA_2_i-NPs and tested their efficacy on cartilage explants from DH guinea pigs. Finally, we investigated the therapeutic effects of sPLA_2_i-NPs in blocking the onset of spontaneous OA in DH guinea pigs. In addition to conventional histologic examination, longitudinal gait analyses were performed to functionally demonstrate improvements of OA-related pain, stiffness, and mobility following sPLA_2_i-NP treatment.

## Material and Methods

### Synthesis of sPLA_2_i-loaded phospholipid NPs

sPLA_2_i-NPs were fabricated via a thin-film hydration approach as previously described. Briefly, lipids composed of 10 mol% 1,2-dioleoyl-3-trimethylammonium-propane (DOTAP, # 890890, Avanti Polar Lipids Inc), a cationic lipid, 25 mol% thioetheramide-PC (#62750, Cayman Chemical), and 65 mol% 1,2-distearoyl-sn-glycero-3-phosphoethanolamine-N-[methoxy(polyethylene glycol)-2000] (DSPE-PEG2000, # 880120, Avanti Polar Lipids Inc), were dissolved in chloroform in a round-bottom flask, with the thioetheramide-PC content fixed at 0.25 mg. Organic solvent was evaporated under a continuous nitrogen flow, followed by vacuum drying for at least 4 h to ensure complete solvent removal. The resulting lipid film was rehydrated with deionized water and incubated at 25 °C for 5 min, after which the suspension was vortexed for an additional 3 min to promote nanoparticle formation.

Large aggregates were removed by centrifugation at 3000 × g for 5 min. The supernatant was subsequently passed through a 0.22 μm cellulose acetate syringe filter (Nalgene, Thermo Scientific), and the purified nanoparticles were stored at 4 °C in the dark until use. Control nanoparticles (Ctrl-NPs) lacking sPLA_2_i with 10 mol% DOTAP, 25 mol% hydrogenated soy phosphatidylcholine (HSPC, # 840058, Avanti Polar Lipids Inc), and 65 mol% DSPE-PEG2000 were prepared following the same procedure. The successful synthesis of NPs was confirmed by dynamic light scattering (DLS; Zetasizer Nano-ZS, Malvern) for size characterization, while the positive surface charge of DOTAP-doped NPs was verified by zeta potential measurement.(7)

For experiments requiring free sPLA_2_i, thioetheramide-PC was initially dissolved in dimethyl sulfoxide (DMSO) to generate a concentrated stock solution, which was then diluted in PBS at a minimum dilution factor of 1:1000 prior to explant and in vivo applications.

### Animal care

In accordance with the standards for animal housing, female DH guinea pigs at 2 months of age were purchased from Charles River and group-housed (2 per cage) at 23-25°C under a 12 h light/dark cycle, with free access to water and standard laboratory pellets. All work performed on animals was approved by the Institutional Animal Care and Use Committee at the University of Pennsylvania.

At 3 months of age, they began to receive treatments once every 2-3 weeks under anesthetic and sterile conditions. To do so, knees were kept in a flexed position and injected with 100 μL of control-NPs in the left knees and sPLA_2_i-NPs (sPLA_2_i: 0.25 mg/mL) in the right knees via an insulin needle. This administration was repeated once every 2-3 weeks until euthanasia

### Micro-computed tomography (microCT) analysis

Bilateral knees were collected and fixed in 4% paraformaldehyde (PFA) for 24 h. Afterwards, they were imaged by micro-computed tomography 45 (µCT45; Scanco Medical AG) at an isotropic voxel resolution of 7.4 µm. Reconstructed images were first processed using a Gaussian smoothing filter (σ = 1.2, support = 2.0) to reduce noise, followed by application of a global threshold set at 30.3% of the maximum grayscale value to segment mineralized tissue from unmineralized or poorly mineralized regions. Sagittal images were contoured for the subchondral bone plate (SBP) followed by generating a 3D color map of thickness for the entire SBP along with a scale bar. This map was then converted to a grayscale thickness map. The region of interest (ROI) was circled from the medial tibial plateau and the average SBP thickness within ROI is calculated by average grey value/255 * max scale bar value. The medial tibial SBP was manually contoured to quantify bone volume fraction (BV/TV), trabecular thickness (Tb.Th), trabecular number (Tb.N), and trabecular separation (Tb.Sp). Coronal images were also contoured followed by 3D reconstruction to count osteophytes at the anterior side.

### Histology

After microCT scanning, knee joints were cut in the middle. The medial halves were decalcified in Morse’s solution (10% sodium citrate and 22.5% formic acid) for 8-10 weeks, followed by tissue processing, paraffin embedding, and sagittal sectioning at 6 μm thickness. Sections with cartilage damage, a total of ∼80-100 per joint, were collected. Every 2 out of 6 consecutive sections were stained by Safranin O/Fast Green (SO/FG) to measure Mankin Score based on OARSI recommendations, (26) which is a sum of scores for cartilage structure (0–8), proteoglycan content (0–6), cellularity (0–3), and tidemark integrity (0–1). The region showing the most severe cartilage degeneration in each section was analyzed, and each knee received a single score representing the maximal score of all its SO/FG-stained sections.

Another 2 out of 6 consecutive sections were stained by hematoxylin and eosin (H&E) to evaluate synovial inflammation using a semi-quantitative grading system in accordance with OARSI histopathology recommendations.(26) Synovial changes including lining layer hyperplasia (0–2), villous hyperplasia (0–3), and degree of cellular infiltration by perivascular lymphocytes and mononuclear cells (0–5) were evaluated and graded on an ordinal scale and summed up for a final score. To ensure consistency, synovitis was quantified at the same location (anterior side, next to femur). Each knee was assigned a single score corresponding to the maximal value observed among its sections.

For immunohistochemical (IHC) analysis, paraffin sections were treated by proteinase K for antigen retrieval and then incubated with primary antibodies overnight at 4 °C. After washing, sections were treated with biotinylated secondary antibodies for 1 h, followed by Elite ABC reagent (Vector Laboratories, PK6100) and 3,3′-diaminobenzidine (DAB; Thermo Scientific, TA-060-QHDX) for visualization. Primary antibodies used in this study included rabbit anti-sPLA_2_IIa antibody (Abcam, ab23705), mouse anti-8-OHdG monoclonal antibody (Abcam, ab48508), rabbit anti-Adamts5 antibody (Abcam, ab41037), and rabbit anti-Mmp13 antibody (AF5355). Quantification was performed by counting the percentage of positively stained cells across the entire articular cartilage. All histological scoring and cell quantification analyses were conducted independently by two blinded evaluators.

To assess systemic toxicity, major organs (heart, lung, liver, spleen, and kidney) were collected, fixed in 4% PFA for 24 h, and processed for paraffin embedding and H&E staining.

### Guinea pig cartilage explant culture

Cartilage explants (2 mm diameter × 2 mm thickness) were harvested from either tibial plateau or femoral condyle of 2.5-month-old DH guinea pigs by biopsy punch. They were first washed in PBS with 100 μg/mL streptomycin and 100 U/mL penicillin for three times and then cultured in chemically defined medium (DMEM, 100 μg/mL streptomycin, 100 U/mL penicillin, 1% ITS+Premix, 40 μg/mL L-proline, 0.1 μM dexamethasone, 100 μg/mL sodium pyruvate, and 50 μg/mL ascorbate 2-phosphate) in a 6-well plate. Three days later, PBS, IL-1β and Ctrl-NP, IL-1β and free sPLA_2_i, or IL-1β and sPLA_2_i-NP were added to the medium to reach final concentrations of IL-1β and sPLA_2_i at 10 ng/mL and 0.1 mg/mL, respectively. Medium was replaced every other day with new treatments. Five days after the initial treatment, explants were collected, fixed with 4% PFA overnight, and processed for paraffin sections with SO/FG staining. OA severity was evaluated by the modified Mankin score for guinea pigs based on the section with the most severe loss of safranin-O staining and cartilage damage.(26) <u>Complete blood count (CBC) analysis</u>

Following euthanasia, whole blood was immediately collected from DH guinea pigs by cardiac puncture using a 1 mL syringe and subjected to CBC analysis using an Element HT5 hematology analyzer (Heska, Loveland, CO).

### sPLA_2_ enzyme activity

A commercial sPLA_2_ assay kit (#765001, Cayman Chemical) was used to evaluate the inhibitory efficiency of sPLA_2_i-NPs and to quantify sPLA_2_ activity in synovial fluid, following the manufacturer’s instructions. Enzymatic activity was determined from the linear region of the absorbance curve at 414 nm. For inhibition analysis, sPLA_2_ activity was measured in the presence of PBS, Ctrl-NPs, free sPLA_2_i, and sPLA_2_i-NPs, and values were normalized to the PBS-treated group (positive control, 100%).

Inhibitory efficiency was calculated as 100% minus the normalized sPLA_2_ activity. Synovial fluid was collected immediately after euthanasia by injecting 100 μL PBS into the closed knee joint cavity using an insulin syringe, followed by aspiration of approximately 100 μL fluid. Samples were stored at −80 °C until analysis.

### Quantitative RT-PCR (qRT-PCR)

Articular cartilage and synovium/infrapatellar fat pad (Syn/IFP) tissue dissected from DH guinea pigs were collected into Tri Reagent (Sigma) and processed for RNA isolation. iScript™ cDNA Synthesis Kit (#1708891, Bio-Rad) was used to reverse-transcribe mRNA into cDNA. Following this, PCR was performed using a Power SYBR Green PCR MasterMix kit (#4367659, Applied BioSystems). The primer sequences for the genes used in this study are listed in Supplementary Table 1.

### Gait analysis

A custom-built transparent acrylic walkway was used as previously reported (Supplementary Figure 1).(27) Specifically, it was 600 mm long and 100 mm wide, with two 130 mm-high acrylic walls at both sides. An entrance and a darkened enclosure were located at two ends to encourage animals to traverse the walkway. Two acrylic force plates (70 mm long, 15 mm thick) were embedded within the walkway surface and positioned 150 mm from the entrance. Beneath them were miniature six-degree-of-freedom force sensors (Nano17; 17 mm diameter; ATI Industrial Automation, Apex, NC). Mirrors were mounted on both sides of the walkway, enabling a digital camera positioned beneath the walkway (A601fc, Basler Vision Technologies, Ahrensburg, Germany) to capture simultaneously ventral and sagittal views of the animal. Image data were collected at 60 frames per second using custom software written in LabVIEW (v8.2, National Instruments, Austin, TX). Kinetic data were subsequently temporally interpolated to match the image acquisition time points using MATLAB (The MathWorks Inc., Natick, MA).(27)

### Statistical analysis

Data are expressed as means ± standard deviation (s.d) and analyzed by paired or unpaired t-tests, one-way ANOVA with Tukey’s post-hoc test and two-way ANOVA with Bonferroni’s or Tukey’s post-hoc test for multiple comparisons using Prism 8 software (GraphPad Software, San Diego, CA). For assays using cartilage explants, experiments were repeated at least three times independently, and representative data from one experiment are shown here. Values of p<0.05 were considered statistically significant.

## Results

### Joint sPLA_2_ increases as DH guinea pigs develop spontaneous OA

To characterize the progression of spontaneous OA in guinea pigs, we first evaluated joint histopathology at 2.5, 6, and 12 months of age using SO/FG and H&E staining. At 2.5-month-old, articular cartilage displayed an intact, smooth surface with preserved structure (Figure 1Aa). In contrast, 6-month-old joints exhibited clear surface degeneration (Figure 1Ab), and by 12 months of age, pronounced cartilage erosion was observed (Figure 1Ac). The corresponding Mankin scores were 0.3, 8.3, and 10.7, respectively (Figure 1B). Synovial pathology progressed in parallel. Compared with the thin synovium in 2.5-month-old joints (Figure 1Ad), the synovium thickened evidently at 6 months of age and further exacerbated at 12 months of age (Figure 1Ae, f), resulting in higher synovitis scores with increasing age (Figure 1B).

**Figure 1.**
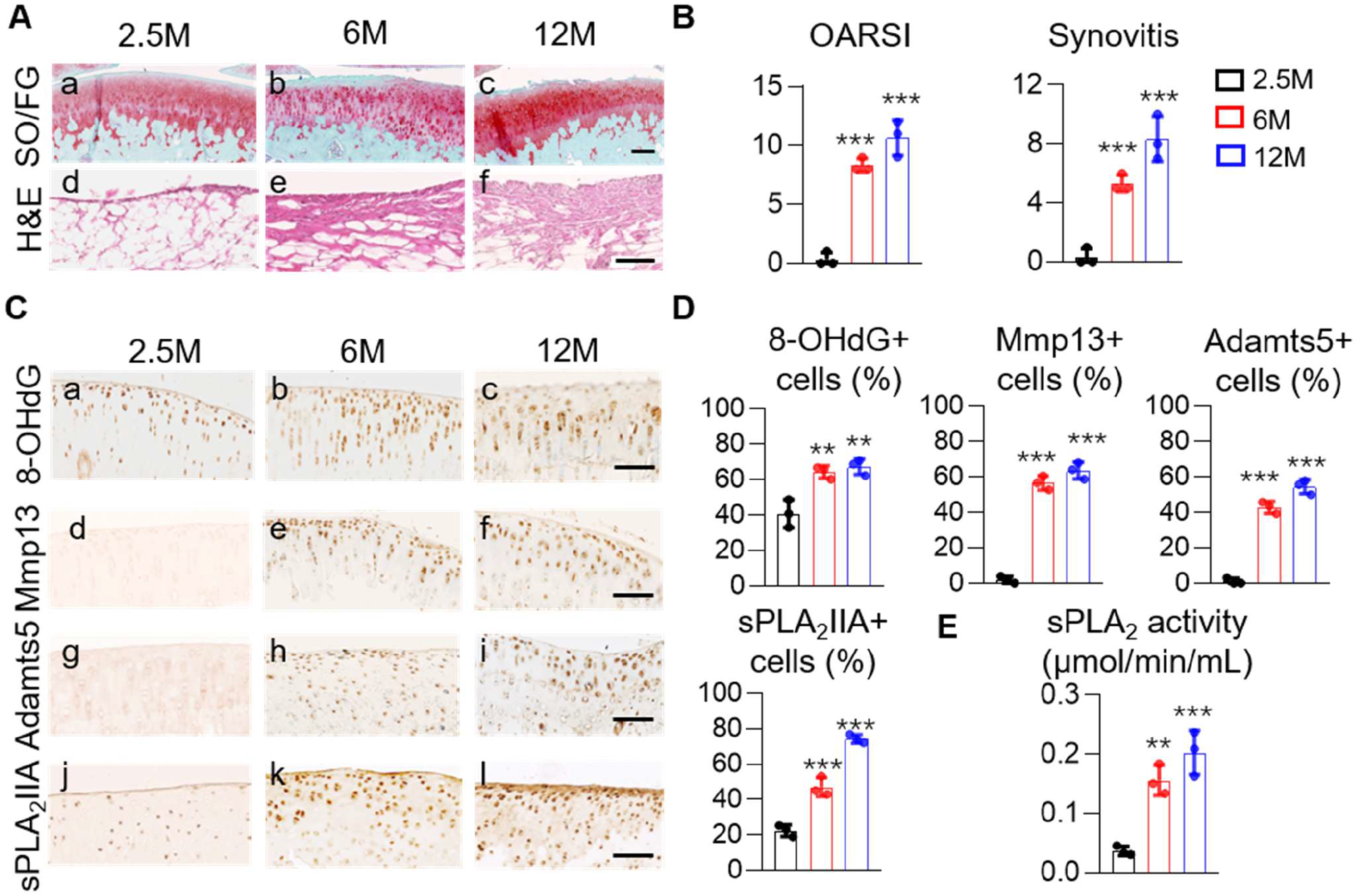
DH guinea pigs develop early-onset spontaneous OA. A, Representative images of tibial articular cartilage (SO/FG staining, a-c) and synovium (H&E staining, d-f) from 2.5-, 6-, and 12-month-old DH guinea pigs. Scale bar, 500μm (a-c) and 100μm (d-f). B, Mankin score for articular cartilage and synovitis score were quantified. (n=3). C, Representative 8-OHdG, Mmp13, Adamts5, and sPLA_2_IIA staining of articular cartilage from DH guinea pigs at different ages. Scale bar, 100μm. D, The percentages of 8-OHdG-positive, Mmp13-positive, and Adamts5-positive chondrocytes were counted in the articular cartilage. (n=3). E, The enzymatic activity of sPLA_2_ in 2.5-, 6-, and 12-month-old guinea pig synovial fluid. (n=3). Statistical analysis was performed using one-way ANOVA with Tukey’s post hoc test. Data presented as mean ± s.d. *, comparison between 2.5-month-old and 6/12-month-old. *p<0.05, **p<0.01, ***p<0.001.

To assess cartilage degeneration at the molecular level, we next stained joints to detect markers associated with OA progression. OA is associated with elevated oxidative stress.(28) 8-hydroxy-2-deoxyguanosine (8-OHdG), a biomarker for measuring oxidative DNA damage, was markedly increased at the cartilage surface in 6-month-old joints and extended into the middle zone chondrocytes by 12-month-old (Figure 1Ca-c, D). Matrix metalloproteinase 13 (Mmp13) and A disintegrin and metalloproteinase with thrombospondin motifs 5 (Adamts5), two proteolytic markers associated with OA-related cartilage degeneration, were significantly elevated by 6-month-old and further increased at 12-month-old, whereas minimal expression was detected in 2.5-month-old joints (Figure 1Cd-f, g-i, D). Together, our histological analyses confirm that DH guinea pigs exhibit spontaneous, progressive OA with increasing age.

sPLA_2_IIA is a common isoform of sPLA_2_ enzyme. We previously reported that its level is drastically elevated in mouse and human articular cartilage after OA initiation.(6) Similarly, sPLA_2_IIA level was minimal in the articular cartilage from 2.5-month-old guinea pigs, while abundant in chondrocytes and cartilage matrix in 6- and 12-month-old guinea pigs (Figure 1Cj-l, D). To further confirm this result, we collected synovial fluid from guinea pigs of different ages to assess sPLA_2_ enzymatic activity. Quantification revealed a significant increase in its enzymatic activity in 6- and 12-month-old animals compared to 2.5-month-old animals (Figure 1E). Together, these results reveal that spontaneous OA in guinea pigs is accompanied by age-related increases in sPLA_2_ expression and activity, indicating that targeting sPLA_2_ could be an efficient approach for treating spontaneous OA in guinea pigs.

### sPLA_2_i-NPs block cartilage degeneration in OA cartilage explants

To inhibit sPLA_2_ activity, we next synthesized NPs encapsulating the sPLA_2_ inhibitor thioetheramide-PC (sPLA_2_i-NPs), along with control nanoparticles (Ctrl-NPs; see Materials and Methods for details, Figure 2A). The NPs were formulated with the cationic lipid DOTAP, resulting in a positively charged surface that may facilitate interactions with negatively charged cellular membranes. Importantly, the lipid shell of the sPLA_2_i-NPs is susceptible to enzymatic degradation by sPLA_2_ present in the environment, enabling stimulus-responsive release of the encapsulated inhibitor. The inhibitory efficiency against sPLA_2_ was assessed using the sPLA_2_ activity kit immediately after synthesis. Both free sPLA_2_i and sPLA_2_i-NPs significantly inhibited enzymatic activity with comparable efficacy, whereas PBS and Ctrl-NPs showed no significant effect (Figure 2B).

**Figure 2.**
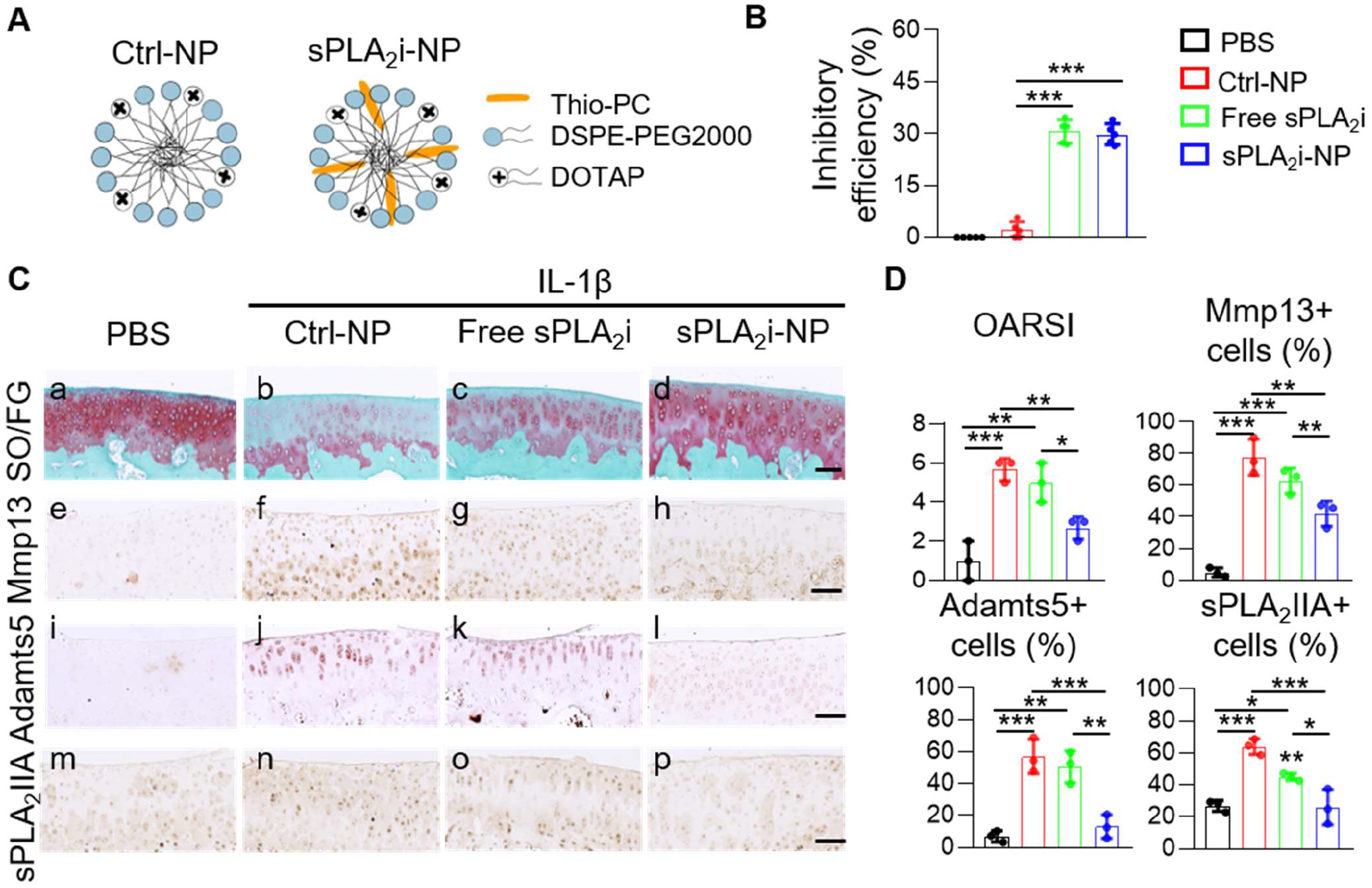
sPLA_2_i-NPs exhibited chondroprotective effects of in cartilage explants from 2.5M Guinea pig. A, Schematic diagram of Ctrl-NPs and sPLA_2_i-NPs. In the sPLA_2_i-NPs, lipid-based sPLA_2_i Thio-PC was incorporated into nanometer-sized phospholipid NPs (DSPE-PEG2000) doped with cationic lipid DOTAP. B, The inhibitory efficiency against sPLA_2_ enzyme of PBS, Ctrl-NP, free sPLA_2_i, and sPLA_2_i-NPs. (n=5). C, Representative images of SO/FG staining(a-d) and IHC staining of Mmp13 (e-h), Adamts5 (i-l), and sPLA_2_IIA (m-p) on the sections of PBS-only, Ctrl-NP-, free sPLA_2_i- and sPLA_2_i-NP-treated IL-1β-stimulated DH guinea pig cartilage explants. Scale bar, 100μm. D, Mankin score and IHC quantification of cartilage explants treated with PBS-only, Ctrl-NP+ IL-1β, free sPLA_2_i+ IL-1β, and sPLA_2_i-NP+IL-1β. (n=3). Statistical analysis was performed using one-way ANOVA with Tukey’s post hoc test. Data presented as mean ± s.d. *p<0.05, **p<0.01, ***p<0.001.

To evaluate their therapeutic effects, we then employed an ex vivo cartilage explant model with a physiologically relevant three-dimensional environment. Healthy articular cartilage explants from 2.5-month-old guinea pigs were treated with interleukin-1β (IL-1β) to induce OA-like phenotypes in the presence of PBS, Ctrl-NPs, free sPLA_2_i, or sPLA_2_i-NPs until day 8. IL-1β stimulation with Ctrl-NPs induced an OA-like phenotype characterized by surface fibrillation and proteoglycan loss (Figure 2Ca, b). Notably, sPLA_2_i-NPs, but not sPLA_2_i, effectively prevented cartilage degeneration, preserving Safranin-O staining intensity, resulting in a Mankin score (2.7) comparable to Ctrl-NP-treated explants (5.7, Figure 2Cb-d, D). The limited efficacy of free sPLA_2_i likely reflects poor penetration of the inhibitor into cartilage in the absence of the positively charged NP carrier. IL-1β together with Ctrl-NPs increased the amounts of catabolic proteases, such as Mmp13 (Figure 2Ce, f) and Adamts5 (Figure 2Ci, j), and sPLA_2_IIA (Figure 2Cm, n) in the cartilage explants. However, sPLA_2_i-NPs, but not free sPLA_2_i, attenuated these increases by 45.9%, 77.3%, 59.3%, respectively (Figure 2Cg, h, k, l, o, p, D).

Collectively, these results demonstrate that sPLA_2_i-NPs can protect the guinea pig cartilage against inflammatory cartilage degeneration and provide ex vivo evidence supporting their therapeutic potential for OA.

### sPLA_2_i-NPs mitigate age-related joint destruction in DH guinea pigs

Next, we examined the therapeutic efficacy of sPLA_2_i-NPs in the OA joints of DH guinea pigs. Starting at 3 months of age, female guinea pigs received intra-articular injections of sPLA_2_i-NP into the right knees and Ctrl-NPs into the left knees once every 2-3 weeks. Animals were sacrificed at 7 and 10 months for histological and microCT analyses.

In line with the above spontaneous OA data, we observed significant cartilage surface damage and synovial thickening in Ctrl-NPs-treated joints of 7-month-old guinea pigs (Figure 3Aa, e), which were further exacerbated in Ctrl-NPs-treated joints of 10-month-old guinea pigs (Figure 3Ac, g). Strikingly, knees receiving 4 months of sPLA_2_i-NP treatment exhibited a relatively smooth, intact cartilage surface and much less synovial swelling (Figure 3Ab, f), resulting 75.0% and 60.0% decreases in Mankin scores and synovitis scores, respectively (Figure 3B). In 10-month-old animals (7 months of sPLA_2_i-NP treatment), cartilage degeneration and synovitis were also significantly attenuated compared with Ctrl-NP-treated knees (Figure 3Ac, d, g, h), with 56.6% and 51.3% decreases in Mankin scores and synovitis scores, respectively (Figure 3B). Meanwhile, IHC staining revealed significant decreases in cartilage sPLA_2_IIA levels after 4 and 7 months of sPLA_2_i-NP treatment (45.4% and 46.1%, respectively, Figure 3C, D).

**Figure 3.**
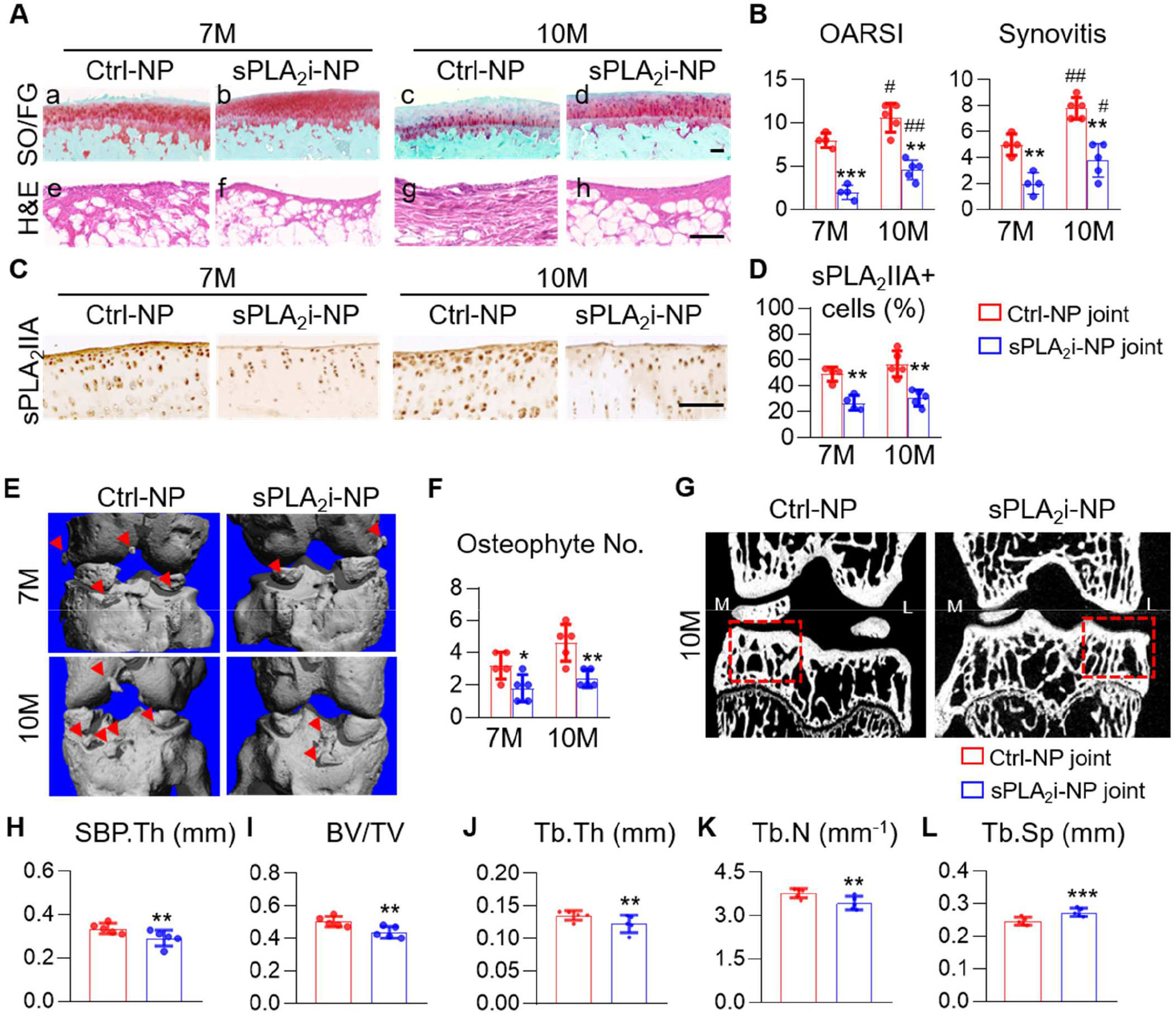
sPLA_2_i-NPs attenuated spontaneous OA progression in DH guinea pig. A, Representative images of SO/FG (a-d) and H&E (e-h) staining of articular cartilage and synovium, respectively, from 7-month- and 10-month-old guinea pigs. Scale bar, 100μm. B, Quantification of Mankin score (left) and synovitis score (right) in healthy and OA guinea pigs. (n=4-5). C, Representative images of IHC staining of sPLA_2_IIA in the tibial articular cartilage. Scale bar, 100μm. D, Quantification of the percentage of sPLA_2_IIA⁺ chondrocytes in total cartilage. (n=4-5). E, Representative images of microCT 3D reconstruction of 7- and 10-month-old guinea pig joints treated with Ctrl-NPs or sPLA_2_i-NPs. Osteophytes on the anterior side were indicated by red arrows. F, Quantification of osteophyte number on the anterior side from the 3D images of 7- and 10-month-old guinea pig joints. (n=5). G, Representative 2D coronal images of the left knee (Ctrl-NP-treated) and right knee (sPLA_2_i-NP-treated) at 10 months of age. The medial subchondral bone region is indicated by red squares. H, Quantification of medial SBP thickness in guinea pig joints at 10 months of age (n=5). I, Quantification of medial subchondral bone volume fraction (BV/TV) in guinea pig joints at 10 months of age (n=5). J, Quantification of trabecular thickness (Tb.Th) in the medial subchondral bone at 10 months of age (n=5). K, Quantification of trabecular number (Tb.N) in the medial subchondral bone at 10 months of age (n=5). L, Quantification of trabecular separation (Tb.Sp) in the medial subchondral bone at 10 months of age (n=5). Statistical analysis was performed using paired t-test between left and right knees, and unpaired t-test between 7- and 10-month-old. Data presented as mean ± s.d. *, comparison between left and right knees; #, comparison between 7-month-old and 10-month-old. *p<0.05, **p<0.01, ***p<0.001; #p<0.05, ##p<0.01.

As late-stage OA is also characterized by subchondral bone remodeling,(29) we further assessed subchondral bone plate (SBP) sclerosis and osteophyte formation. Compared to Ctrl-NP treatment, sPLA_2_i-NP treatment significantly reduced osteophytes by 43.8% and 47.8% in 7- and 10-month-old guinea pigs (Figure 3E, F). Consistently, sPLA_2_i-NP treatment attenuated pathological thickening of the medial SBP by 13.3% (Figure 3H), reduced medial subchondral bone volume fraction (BV/TV) by 12.9% (Figure 3I), and decreased trabecular thickness (Tb.Th) and trabecular number (Tb.N) of the medial subchondral bone by 9.5% and 9.1%, respectively (Figure 3J, K). In contrast, medial trabecular separation (Tb.Sp) increased by 11.3% (Figure 3L). Together, these findings demonstrate that sPLA_2_i-NPs not only preserve cartilage integrity but also suppress synovial inflammation and pathological bone remodeling, effectively delaying the progression of spontaneous OA.

To further characterize the effect of sPLA_2_i-NPs on cartilage metabolism, we measured the expression of anabolic and catabolic genes in articular cartilage and synovium/infrapatellar fat pad (Syn/IFP) collected from bilateral knees of 10-month-old guinea pigs. We also collected samples from 2.5-month-old guinea pigs with no injections as healthy controls. Consistent with spontaneous OA development in guinea pigs, qRT-PCR analysis revealed decreased *Col2a1* expression (46.8%, anabolic) and increased the expression of catabolic (*Mmp13*, 3.6-fold; *Adamts5*, 3.4-fold) and inflammatory cytokine (*Il-1b*, 4.1-fold) genes in 10-month-old Ctrl-NPs-treated cartilage compared to 2.5-month-old healthy samples. Notably, the expression of *Pla2g2a*, the gene encoding sPLA_2_IIA, increased 1.9-fold, consistent with its elevated protein amount (Figure 4A). In the 10-month-old samples, sPLA_2_i-NP treatment greatly increased *Col2a1* (1.3-fold) expression and decreased *Mmp13* (50.1%), *Adamts5* (54.7%), *Il-1b* (58.9%), and *Pla2g2a* (31.0%). qRT-PCR assays on synovium and fat pad issues revealed similar changes in catabolic and inflammatory genes (Figure 4B). To further confirm the reduction of *Pla2g2a* gene expression after sPLA_2_i-NP treatment, we measured sPLA_2_ activity in synovial fluid from 10-month-old animals. As expected, sPLA_2_i-NP-treated knees exhibited significantly reduced enzymatic activity compared to Ctrl-NP-treated knees (Figure 4C). Taken together, our data demonstrates the in vivo efficacy of sPLA_2_i-NPs in mitigating spontaneous OA and reveals their actions in regulating joint tissue metabolism.

**Figure 4.**
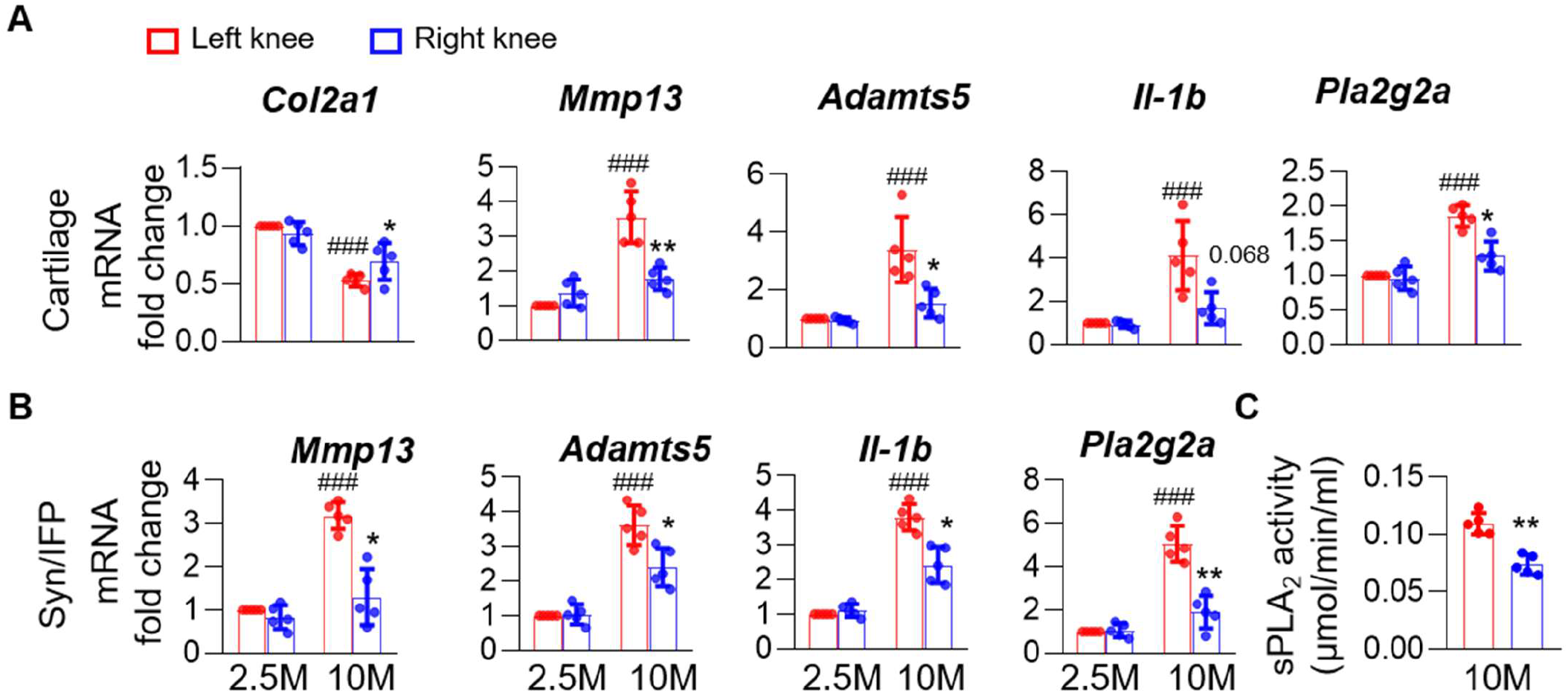
sPLA_2_i-NPs decreased catabolic marker and sPLA_2_ levels while increasing anabolic gene expression. A, qRT-PCR analysis of *Col2a1* (anabolic), *Mmp13*, *Adamts5*, *Il-1b* (catabolic), and *Pla2g2a* expression in the articular cartilage. (n=5). B, qRT-PCR analysis of *Mmp13*, *Adamts5*, *Il-1b* (catabolic), and *Pla2g2a* expression in the synovium. (n=5). C, Enzymatic activity test of sPLA_2_ in the synovial fluid of the left and right knees of 10-month-old DH guinea pigs. (n=5). Statistical analysis was performed using paired t-test between left and right knees, and unpaired t-test between 2.5- and 10-month-old. Data presented as mean ± s.d. *, comparison between left and right knees; #, comparison between 2.5-month-old and 10-month-old. *p<0.05, **p<0.01, ***p<0.001; ###p<0.001.

### sPLA_2_i-NPs maintain normal gait in DH guinea pigs

Gait abnormality is a hallmark of late-stage OA in human patients and reflects their impaired joint function and pain-related weight offloading.(30) To examine the clinical benefits of sPLA_2_i-NP treatment, we next assessed its functional outcomes in 5-, 7-, and 10-month-old guinea pigs that had received 2, 4, and 7 months of sPLA_2_i-NP injections, respectively, using a customized walkway equipped with force plates and high-speed cameras (Figure 5A, Supplementary video 1). This walkway allows us to characterize each leg’s gait separately. 2.5-month-old guinea pigs were used as healthy controls and showed no difference in gait analysis between the two legs. We first measured the locomotor activity (Figure 5B, C). While stride length remained unchanged, stride speed was reduced in the left (untreated or Ctrl-NP-treated) limbs at 7 and 10 months of age. In contrast, the right (sPLA_2_i-NP-treated) limbs exhibited significant increases in both stride length (15.3%) and stride speed (27.7%) in the 7-month-old group compared with their contralateral limbs. These increases were further exaggerated in the 10-month-old group (23.0% in stride length and 71.0% in stride speed), indicating improved locomotor performance.

**Figure 5.**
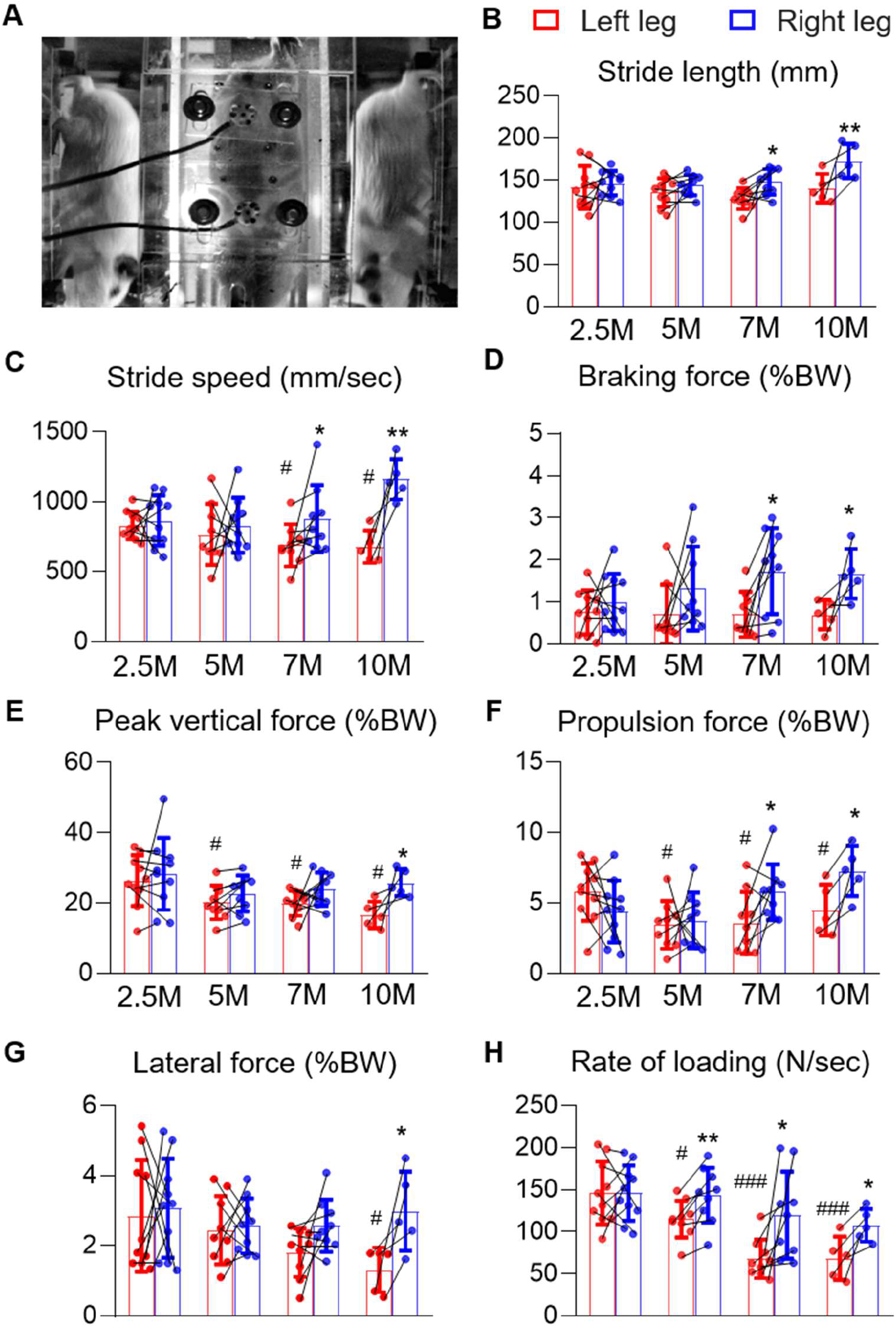
sPLA_2_i-NP treatment ameliorated the OA-related gait abnormality in DH guinea pigs. A, Representative photo of a DH guinea pig walks through the acrylic walkway. B, Quantification of Stride length (mm) of the left (Ctrl-NP-treated) and the right (sPLA_2_i-NP-treated) legs in 2.5-, 5-, 7-, and 10-month-old guinea pigs. C, Quantification of Stride speed (mm/sec) of the left and right legs in 2.5-, 5-, 7-, and 10-month-old guinea pigs. D, Quantification of braking force normalized to body weight (BW) of the left and right legs in 2.5-, 5-, 7-, and 10-month-old guinea pigs. E, Quantification of peak lateral force normalized to BW of the left and right legs in 2.5-, 5-, 7-, and 10-month-old guinea pigs. F, Quantification of propulsion force normalized to BW of the left and right legs in 2.5-, 5-, 7-, and 10-month-old guinea pigs. G, Quantification of lateral force normalized to BW of the left and right legs in 2.5-, 5-, 7-, and 10-month-old guinea pigs. H, Quantification of rate of loading (N/sec) of the left and right legs in 2.5-, 5-, 7-, and 10-month-old guinea pigs. (n=10 for 2.5-month-old animal, n=9 for 5- and 7-month-old animal, and n=5 for 10-month-old animal for panels B-H). Statistical analysis was performed using paired t-test between left and right legs, and unpaired t-test between 2.5-month-old and 5/7/10-month-old. Data presented as mean ± s.d. *, comparison between left and right knees; #, comparison between 2.5-month-old and 5/7/10-month-old. *p<0.05, **p<0.01; #p<0.05, ###p<0.001

This walkway also measures loading and ground reaction force parameters, including braking force, peak vertical force, propulsion force, and lateral force, which together describe weight acceptance, support, push-off, and stability during stance.(31) Across these metrics, left legs exhibited a trend of decline in limb loading during aging, reflected by reduced peak vertical force, propulsion force, and lateral force over time (Figure 5D-G). In contrast, sPLA_2_i-NP-treated right legs consistently bore greater forces than Ctrl-NP-treated left legs. Specifically, compared to left legs, braking force and propulsion force of right legs were significantly increased at both 7 and 10 months of age, while peak vertical force and lateral force were significantly increased at 10 months of age. The rate of loading, which reflects how rapidly body weight is transferred onto the limb, gradually decreased with age in control legs (Figure 5H). Starting from 5 months of age, sPLA_2_i-NP-treated legs showed a significantly improved rate of loading compared to control legs, suggesting earlier recovery of confident limb use. Together, these findings demonstrate that sPLA_2_i-NP treatment restores gait mechanics and weight-bearing capacity, providing functional evidence of disease modification in vivo.

### Long-term administration of sPLA_2_i-NPs does not cause obvious side effects

To evaluate the safety profile of sPLA_2_i-NPs, we assessed potential systemic toxicity following serial intra-articular injections in guinea pigs. A complete blood count (CBC) was performed in 10-month-old animals after 7 months of sPLA_2_i injections and 10-month-old animals with no treatment as controls. No significant differences were observed in total white and red blood cell counts, as well as specific types of blood cells (neutrophils, eosinophils, lymphocytes, monocytes, and basophils), indicating no detectable hematologic abnormalities or systemic inflammatory responses (Figure 6A).

**Figure 6.**
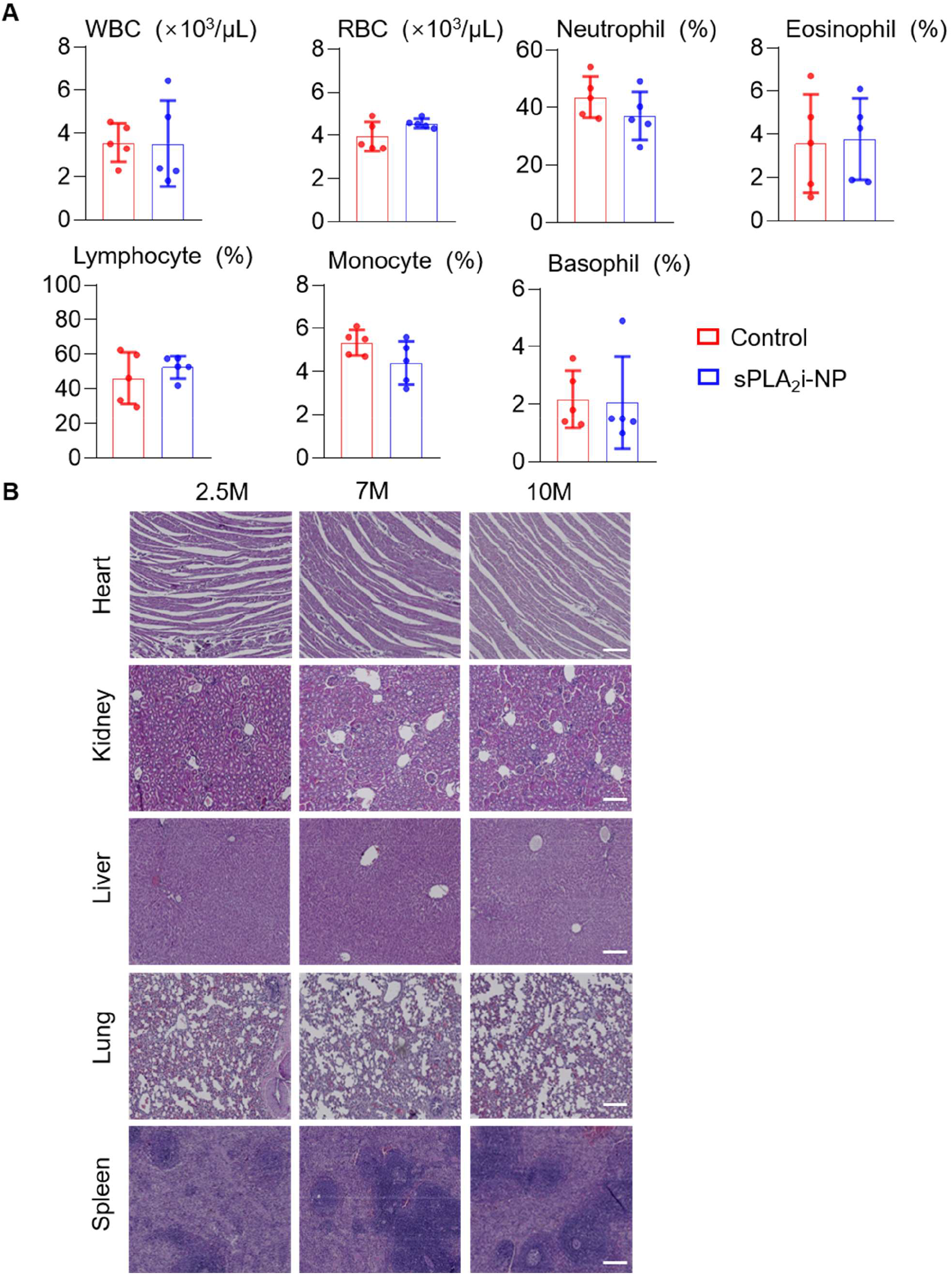
Long-term sPLA_2_i-NP treatment did not result in systemic toxicity. A, CBC blood test of blood collected from 10-month-old DH guinea pigs. WBC, white blood cell; RBC, red blood cell; NEU, neutrophil; EOS, eosinophils; LYM, lymphocyte; MONO, monocyte; BAS, basophils. (n=5). B, Representative images of H&E staining of major organs (heart, kidney, liver, lung, and spleen) collected from non-treated and sPLA_2_i-NP-treated guinea pigs. Scale bar, 200 μm. Statistical analysis was performed using unpaired Student t-test. Data presented as mean ± s.d.

To further assess potential organ toxicity, histopathological analysis was conducted on major organs in 7- and 10-month-old animals after 4 and 7 months of sPLA_2_i-NP treatment. Compared to 2.5-month-old guinea pigs without treatment, these animals showed no histological abnormalities in liver, spleen, kidney, lung, and heart (Figure 6B). Collectively, these findings demonstrate that long-term intra-articular administration of sPLA_2_i-NPs at the tested dosage does not induce detectable systemic or organ toxicity.

## Discussion

In the present study, we demonstrate that targeting sPLA_2_ signaling with advanced NP technology is a promising therapeutic approach for spontaneous, age-related OA and its associated gait abnormality. In our hands, female DH guinea pigs exhibited notable OA symptoms at 5-6 months of age, including cartilage degeneration, synovitis, and reduced ground reaction forces, accompanied by elevated expression and enzymatic activity of sPLA_2_ within the joints. Using this animal model, we found that ex vivo, adding sPLA_2_i-NPs to culture medium protects guinea pig articular cartilage from IL-1β-induced OA degeneration, and that in vivo, administering sPLA_2_i-NPs into guinea pig knees over 4-7 months attenuates cartilage destruction, synovial inflammation, and osteophyte formation, which were further translated into a preservation of gait mechanics. Importantly, we did not detect any obvious, global side effects in the animals. Together, our findings establish sPLA_2_ as a key inflammatory mediator in spontaneous, age-related OA and provide strong preclinical evidence supporting NP-mediated sPLA_2_ inhibition as a clinically translatable disease-modifying strategy.

Accumulating evidence supports the concept that low-grade inflammation is a central driver of OA progression rather than a mere secondary consequence of cartilage breakdown.(5, 32, 33) sPLA_2_ enzymes catalyze the hydrolysis of membrane phospholipids, releasing arachidonic acid and lysophospholipids, which serve as precursors for prostaglandins, leukotrienes, and other inflammatory mediators.(34) Elevated sPLA_2_ expression and activity have been reported in human OA synovium and cartilage,(6, 10, 12) yet the causal contribution of sPLA_2_ to OA progression has remained incompletely defined. Our data extend these observations by demonstrating age-dependent increases in sPLA_2_-IIA expression and synovial fluid enzymatic activity during spontaneous OA development in DH guinea pigs. These findings are consistent with our previous mouse and human studies showing robust sPLA_2_ upregulation during OA initiation and progression. Importantly, inhibition of sPLA_2_ not only reduced inflammatory gene expression (*Il-1b*, *Pla2g2a*) but also suppressed catabolic mediators (*Mmp13*, *Adamts5*) while restoring anabolic gene expression (*Col2a1*), indicating that sPLA_2_ sits upstream of multiple destructive pathways. Compared with single-cytokine blockade strategies such as anti-TNF or anti-IL-1β, which have shown limited efficacy in OA,(35, 36) targeting sPLA_2_ may provide broader suppression of inflammatory lipid mediators and downstream catabolic cascades.

Most preclinical OA studies rely on surgically induced models such as DMM, which resemble post-traumatic OA rather than idiopathic, age-related disease.(37, 38) In contrast, DH guinea pigs develop spontaneous, bilaterally symmetric OA that closely mirrors human age-associated OA in terms of medial compartment predominance, progressive cartilage erosion, synovitis, osteophyte formation, and subchondral bone remodeling.(18, 22, 26) In this context, our study is particularly important because it demonstrates therapeutic efficacy in a naturally occurring OA model without surgical induction. The observed age-dependent increase in oxidative stress (8-OHdG) further align with prior reports that oxidative damage contribute to cartilage degeneration.(39) Therefore, our data provide translationally relevant evidence that sPLA_2_ inhibition can modify disease progression in a model that closely recapitulates human spontaneous OA.

One major barrier to intra-articular drug therapy is rapid clearance from the joint space through synovial vasculature and lymphatics.(40) Previous clinical trials of small-molecule inhibitors in inflammatory diseases demonstrated only transient benefits, likely due to insufficient joint retention.(41) Another key limitation is the restricted penetration of free drugs into cartilage tissue. In line with this, free sPLA_2_i failed to protect cartilage in explant cultures, whereas sPLA_2_i-NPs significantly preserved proteoglycan content and structural integrity. Cartilage contains a dense, negatively charged extracellular matrix rich in glycosaminoglycans, which can be leveraged for electrostatic-based targeting using cationic carriers.(42–44) Our DOTAP-doped phospholipid micelles exhibited slightly positive surface charge, enabling enhanced cartilage penetration and retention while maintaining favorable biocompatibility. The functional superiority of sPLA_2_i-NPs over free inhibitor underscores the critical importance of drug-delivery design for achieving disease modification in OA.

Various nanoparticle systems, including liposomes, dendrimers, and polymeric particles, have been explored for OA therapy.(40) While dendrimers offer excellent cartilage penetration,(45, 46) their clinical translation has been limited. Liposomes, although biocompatible, may not reach deep cartilage layers efficiently.(40) In contrast, our phospholipid micelles consist largely of clinically utilized lipid components and allow straightforward incorporation of lipid-based inhibitors without chemical conjugation. Moreover, the sPLA_2_-responsive design potentially enables enzyme-triggered therapeutic action in inflamed joints. Thus, this platform combines translational feasibility with mechanistic targeting.

Structural preservation does not necessarily translate into functional benefit. Therefore, an important strength of this study is the incorporation of longitudinal gait analysis.(47) Gait abnormalities in knee OA patients reflect pain, impaired weight acceptance, reduced propulsion, and instability.(30) Ground reaction force parameters, including braking force, peak vertical force, propulsion, lateral force, and rate of loading, quantify these biomechanical deficits.(31) Using a custom-built walkway, we observed progressive age-related declines in limb loading parameters in control-NP-treated limbs, which is consistent with worsening spontaneous OA. In contrast, sPLA_2_i-NP-treated limbs maintained higher propulsion, peak vertical force, and rate of loading, suggesting improved weight-bearing confidence and joint function. These functional improvements paralleled reductions in cartilage degeneration, synovitis, and osteophyte formation, supporting the concept that sPLA_2_ inhibition produces true disease modification rather than merely histological improvement.

Several limitations should be acknowledged. First, only female animals were used in the current study. Previous work by Wang et al. demonstrated that male DH guinea pigs exhibited more severe cartilage degeneration than females, likely due to their faster growth rate and greater body weight.

However, both sexes displayed similar age-dependent trends in OA progression and subchondral bone remodeling. (48) Second, the sample size in this study was limited, which may have affected the statistical power of the behavioral analyses. For example, no significant changes in braking force were observed in the left limb (Ctrl-NP-treated) with aging, and lateral force reached significance only at 10 months. Nevertheless, a previous study in 2012 reported that braking and mediolateral forces did not change significantly in a rat OA model following medial meniscus transection, suggesting that these gait parameters may not be consistently sensitive indicators of OA progression.(49) Finally, thioetheramide-PC is a research-grade inhibitor with moderate potency; clinically validated sPLA_2_ inhibitors with lower IC₅₀ values may further enhance therapeutic efficacy.

In summary, this study demonstrates that sPLA_2_ is progressively upregulated during spontaneous OA development in DH guinea pigs and that sustained intra-articular delivery of sPLA_2_i-NPs significantly attenuates cartilage degeneration, synovitis, pathological bone remodeling, and gait dysfunction. By targeting an upstream inflammatory enzyme and overcoming intra-articular delivery barriers through nanoparticle engineering, our strategy provides compelling preclinical evidence for sPLA_2_ inhibition as a disease-modifying therapy for age-related OA. Future studies should evaluate therapeutic efficacy in established OA, optimize nanoparticle formulation for larger animal models, and incorporate clinically advanced sPLA_2_ inhibitors to facilitate translation. Given the persistent unmet need for disease-modifying OA drugs, nanoparticle-mediated targeting of sPLA_2_ represents a promising and mechanistically grounded therapeutic avenue.

## Acknowledgments

We thank Drs. Victoria Lynn Anderson, Jennifer M Kirsch, James Olmsted Marx, Leah Makaron, Kristen Esannason Munroe, Charissa T Wiley, Alessandro P Lamacchia, Caitlin Homanick, Kristin Lee Gardiner, Imani Nicolis, Stuart Isaacs, Blythe Philips, Derrick A Dow, Joseph M Altemus, and Ariel Ayers at Penn University Laboratory Animal Resources (ULAR) for assisting us with guinea pig maintenance, intra-articular injection, and euthanasia. We thank Dr. Louis Soslowsky for offering the custom-built transparent acrylic walkway for gait analysis. This study was supported by NIH grants R01AG067698 (to L.Q.), P30AR069619 (to Penn Center for Musculoskeletal Disorders), R01AR080820 (to Z.C.), R01AR086381 (to Z.C.), and the Health Research Formula Fund from the Pennsylvania Department of Health (to Z.C.).

---

**Supplementary Figure 1.**
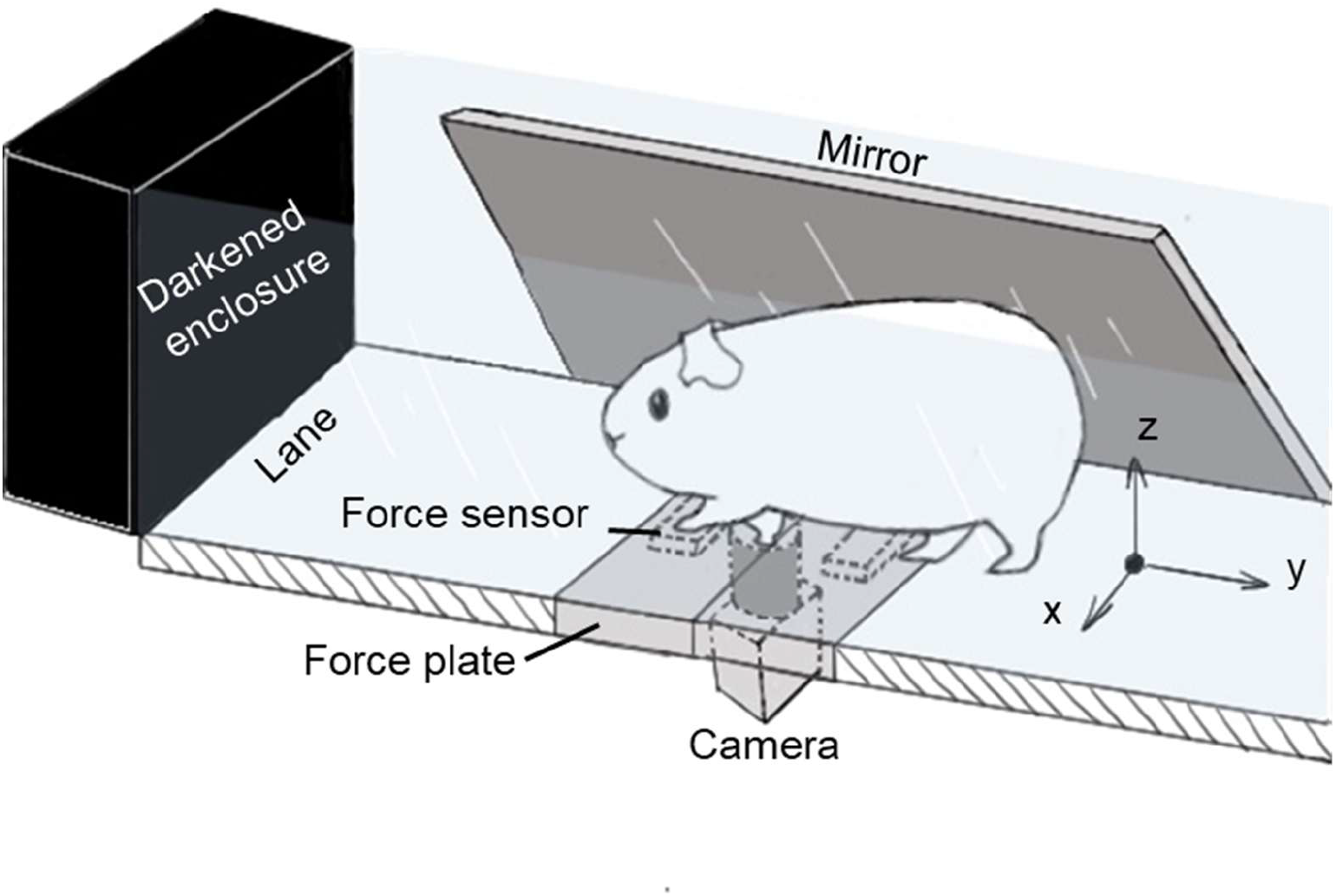
Schematic diagram of a custom-made walkway for rodent gait analysis. To encourage forward locomotion, a darkened enclosure was positioned 600 mm beyond the exit of a custom-built transparent acrylic walkway. The walkway (15 mm thick, 100 mm wide) was bounded by two clear acrylic sidewalls (130 mm high). Ground reaction forces were recorded using two acrylic force plates (70 mm long, 15 mm thick), each instrumented with a miniature six-degree-of-freedom force/torque sensor. Mirrors mounted along both sides of the walkway enabled a digital camera placed beneath the transparent surface to simultaneously capture ventral and bilateral sagittal views of the animal. Figure only shows one of the mirrors.

**Supplementary Video 1.**
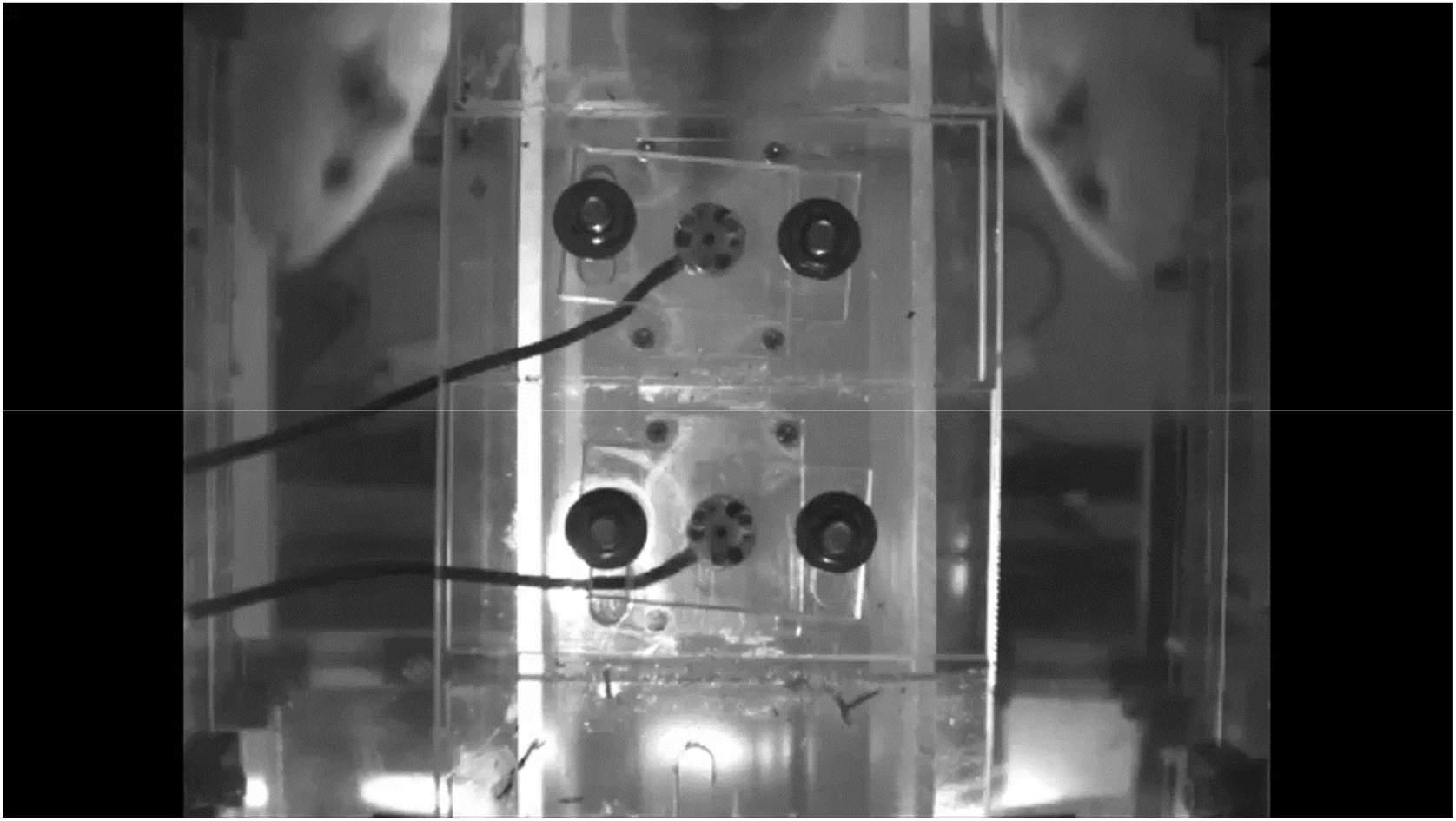
Representative video showing a 7-month-old DH guinea pig walking through the custom-made walkway during one measurement of gait analysis.

**Table 1.**
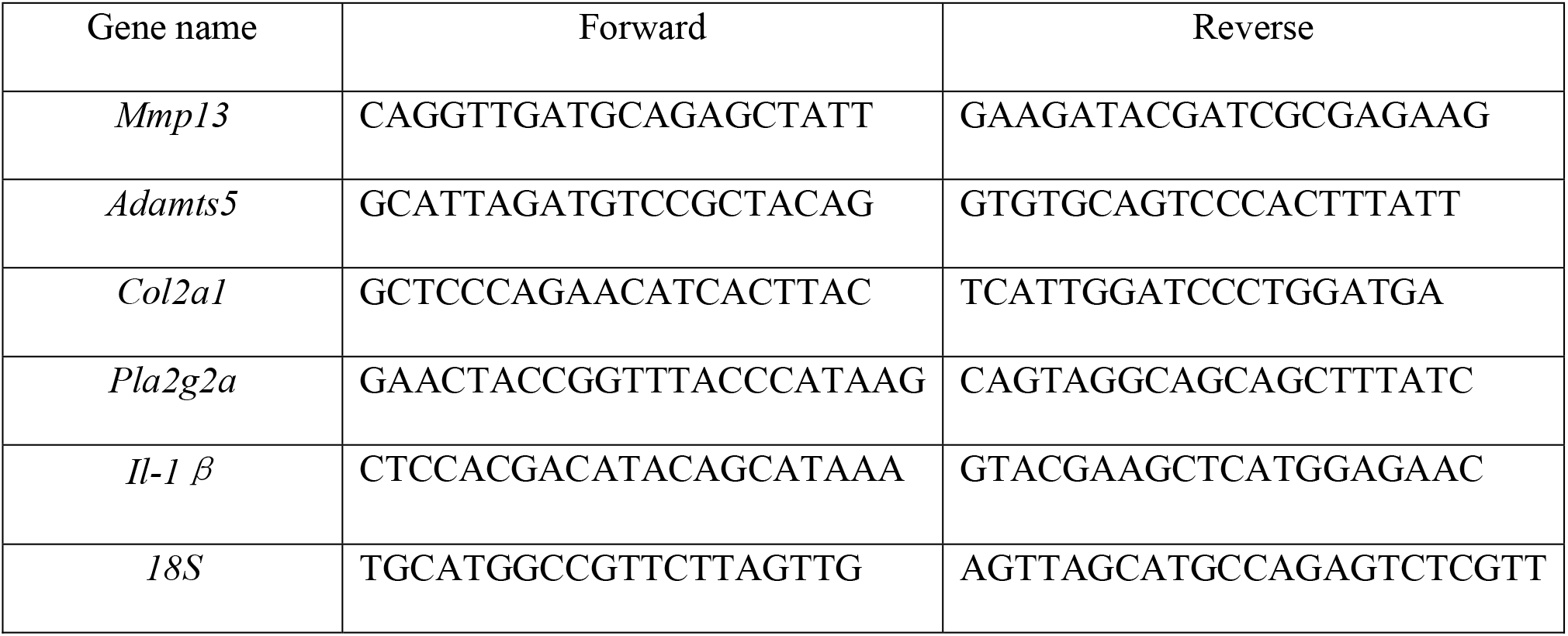
Primer sequences for qRT-PCR analysis in guinea pigs.

## Notes

### Competing Interest Statement

The authors have declared no competing interest.

## References

1. Sharma L. Osteoarthritis of the Knee. The New England journal of medicine. 2021;384(1):51–9.

2. Johnson VL, Hunter DJ. The epidemiology of osteoarthritis. Best Pract Res Clin Rheumatol. 2014;28(1):5–15.

3. Sokolove J, Lepus CM. Role of inflammation in the pathogenesis of osteoarthritis: latest findings and interpretations. Ther Adv Musculoskelet Dis. 2013;5(2):77–94.

4. Lapane KL, Yang S, Driban JB, Liu SH, Dubé CE, McAlindon TE, et al. Effects of prescription nonsteroidal antiinflammatory drugs on symptoms and disease progression among patients with knee osteoarthritis. Arthritis Rheumatol. 2015;67(3):724–32.

5. Robinson WH, Lepus CM, Wang Q, Raghu H, Mao R, Lindstrom TM, et al. Low-grade inflammation as a key mediator of the pathogenesis of osteoarthritis. Nat Rev Rheumatol. 2016;12(10):580–92.

6. Wei Y, Yan L, Luo L, Gui T, Jang B, Amirshaghaghi A, et al. Phospholipase A(2) inhibitor-loaded micellar nanoparticles attenuate inflammation and mitigate osteoarthritis progression. Sci Adv. 2021;7(15).

7. Liang Q, Gui T, Luo L, Popov AV, Isuri RK, He Q, et al. Varespladib-Based Lipid Nanoparticles as Highly Efficient Anti-Inflammatory Agents for Osteoarthritis Treatment. ACS Appl Mater Interfaces. 2025;17(45):61843–54.

8. Bradley JD, Dmitrienko AA, Kivitz AJ, Gluck OS, Weaver AL, Wiesenhutter C, et al. A randomized, double-blinded, placebo-controlled clinical trial of LY333013, a selective inhibitor of group II secretory phospholipase A2, in the treatment of rheumatoid arthritis. J Rheumatol. 2005;32(3):417–23.

9. Zhao Z, Liu N, Huang J, Lu PH, Xu XM. Inhibition of cPLA2 activation by Ginkgo biloba extract protects spinal cord neurons from glutamate excitotoxicity and oxidative stress-induced cell death. J Neurochem. 2011;116(6):1057–65.

10. Jamal OS, Conaghan PG, Cunningham AM, Brooks PM, Munro VF, Scott KF. Increased expression of human type IIa secretory phospholipase A2 antigen in arthritic synovium. Ann Rheum Dis. 1998;57(9):550–8.

11. Pruzanski W, Bogoch E, Wloch M, Vadas P. The role of phospholipase A2 in the physiopathology of osteoarthritis. J Rheumatol Suppl. 1991;27:117–9.

12. Pruzanski W, Bogoch E, Stefanski E, Wloch M, Vadas P. Enzymatic activity and distribution of phospholipase A2 in human cartilage. Life Sci. 1991;48(25):2457–62.

13. Zhang W, Sun G, Aitken D, Likhodii S, Liu M, Martin G, et al. Lysophosphatidylcholines to phosphatidylcholines ratio predicts advanced knee osteoarthritis. Rheumatology (Oxford). 2016;55(9):1566–74.

14. Geraghty T, Obeidat AM, Ishihara S, Wood MJ, Li J, Lopes EBP, et al. Age-Associated Changes in Knee Osteoarthritis, Pain-Related Behaviors, and Dorsal Root Ganglia Immunophenotyping of Male and Female Mice. Arthritis Rheumatol. 2023;75(10):1770–80.

15. Ariosa-Morejon Y, Santos A, Fischer R, Davis S, Charles P, Thakker R, et al. Age-dependent changes in protein incorporation into collagen-rich tissues of mice by in vivo pulsed SILAC labelling. Elife. 2021;10.

16. McNulty MA, Loeser RF, Davey C, Callahan MF, Ferguson CM, Carlson CS. Histopathology of naturally occurring and surgically induced osteoarthritis in mice. Osteoarthritis Cartilage. 2012;20(8):949–56.

17. McCoy AM. Animal Models of Osteoarthritis: Comparisons and Key Considerations. Vet Pathol. 2015;52(5):803–18.

18. Veronesi F, Salamanna F, Martini L, Fini M. Naturally Occurring Osteoarthritis Features and Treatments: Systematic Review on the Aged Guinea Pig Model. International Journal of Molecular Sciences. 2022;23(13):7309.

19. Huebner JL, Otterness IG, Freund EM, Caterson B, Kraus VB. Collagenase 1 and collagenase 3 expression in a guinea pig model of osteoarthritis. Arthritis Rheum. 1998;41(5):877–90.

20. Santangelo KS, Nuovo GJ, Bertone AL. In vivo reduction or blockade of interleukin-1β in primary osteoarthritis influences expression of mediators implicated in pathogenesis. Osteoarthritis and cartilage. 2012;20(12):1610–8.

21. Santangelo KS, Pieczarka EM, Nuovo GJ, Weisbrode SE, Bertone AL. Temporal expression and tissue distribution of interleukin-1β in two strains of guinea pigs with varying propensity for spontaneous knee osteoarthritis. Osteoarthritis and cartilage. 2011;19(4):439–48.

22. Meacock SC, Bodmer JL, Billingham ME. Experimental osteoarthritis in guinea-pigs. J Exp Pathol (Oxford). 1990;71(2):279–93.

23. Gregory MH, Capito N, Kuroki K, Stoker AM, Cook JL, Sherman SL. A review of translational animal models for knee osteoarthritis. Arthritis. 2012;2012:764621.

24. Sato M, Uchida K, Nakajima H, Miyazaki T, Guerrero AR, Watanabe S, et al. Direct transplantation of mesenchymal stem cells into the knee joints of Hartley strain guinea pigs with spontaneous osteoarthritis. Arthritis research & therapy. 2012;14(1):R31.

25. Kwapisz A, Bowman M, Walters J, Cosh H, Burnikel B, Tokish J, et al. Human Adipose- and Amnion-Derived Mesenchymal Stromal Cells Similarly Mitigate Osteoarthritis Progression in the Dunkin Hartley Guinea Pig. Am J Sports Med. 2022;50(14):3963–73.

26. Kraus VB, Huebner JL, DeGroot J, Bendele A. The OARSI histopathology initiative - recommendations for histological assessments of osteoarthritis in the guinea pig. Osteoarthritis Cartilage. 2010;18 Suppl 3(Suppl 3):S35–52.

27. Sarver JJ, Dishowitz MI, Kim SY, Soslowsky LJ. Transient decreases in forelimb gait and ground reaction forces following rotator cuff injury and repair in a rat model. J Biomech. 2010;43(4):778–82.

28. Zahan OM, Serban O, Gherman C, Fodor D. The evaluation of oxidative stress in osteoarthritis. Med Pharm Rep. 2020;93(1):12–22.

29. Burr DB, Gallant MA. Bone remodelling in osteoarthritis. Nature Reviews Rheumatology. 2012;8(11):665–73.

30. Favre J, Jolles BM. Gait analysis of patients with knee osteoarthritis highlights a pathological mechanical pathway and provides a basis for therapeutic interventions. EFORT Open Rev. 2016;1(10):368–74.

31. Vaverka F, Elfmark M, Svoboda Z, Janura M. System of gait analysis based on ground reaction force assessment. Acta Gymnica. 2015;45(4):187–93.

32. Scanzello CR. Role of low-grade inflammation in osteoarthritis. Curr Opin Rheumatol. 2017;29(1):79–85.

33. Terkawi MA, Ebata T, Yokota S, Takahashi D, Endo T, Matsumae G, et al. Low-Grade Inflammation in the Pathogenesis of Osteoarthritis: Cellular and Molecular Mechanisms and Strategies for Future Therapeutic Intervention. Biomedicines. 2022;10(5).

34. Kudo I, Murakami M. Phospholipase A2 enzymes. Prostaglandins & Other Lipid Mediators. 2002;68–69:3–58.

35. Dimitroulas T, Lambe T, Klocke R, Kitas GD, Duarte RV. Biologic drugs as analgesics for the management of osteoarthritis. Seminars in Arthritis and Rheumatism. 2017;46(6):687–91.

36. Knapik M, Żelazo DA, Osowiecka K, Krajewska-Włodarczyk M. Efficacy of Anti-Interleukin-1 Therapeutics in the Treatment of Knee Osteoarthritis: A Systematic Review and Meta-Analysis of Randomized Controlled Trials from the Years 2000 to 2023. J Clin Med. 2024;13(10).

37. Kuyinu EL, Narayanan G, Nair LS, Laurencin CT. Animal models of osteoarthritis: classification, update, and measurement of outcomes. J Orthop Surg Res. 2016;11:19.

38. Teeple E, Jay GD, Elsaid KA, Fleming BC. Animal models of osteoarthritis: challenges of model selection and analysis. Aaps j. 2013;15(2):438–46.

39. Martin JA, Klingelhutz AJ, Moussavi-Harami F, Buckwalter JA. Effects of Oxidative Damage and Telomerase Activity on Human Articular Cartilage Chondrocyte Senescence. The Journals of Gerontology: Series A. 2004;59(4):B324–B36.

40. Liang Q, Cheng Z, Qin L. Advanced nanoparticles in osteoarthritis treatment. Biomater Transl. 2024;5(2):95–113.

41. Brandt MD, Malone JB, Kean TJ. Advances and Challenges in the Pursuit of Disease-Modifying Osteoarthritis Drugs: A Review of 2010-2024 Clinical Trials. Biomedicines. 2025;13(2).

42. Zhang C, Vedadghavami A, He T, Charles JF, Bajpayee AG. Cationic Carrier Mediated Delivery of Anionic Contrast Agents in Low Doses Enable Enhanced Computed Tomography Imaging of Cartilage for Early Osteoarthritis Diagnosis. ACS Nano. 2023;17(7):6649–63.

43. Pathrikar TV, Baby HM, Hakim B, Zhang H, Millán Cotto HA, Kondiboyina V, et al. Cartilage-targeting exosomes for delivery of receptor antagonist of interleukin-1 in osteoarthritis treatment. Osteoarthritis and Cartilage. 2025;33(7):835–47.

44. Bajpayee AG, Grodzinsky AJ. Cartilage-targeting drug delivery: can electrostatic interactions help? Nat Rev Rheumatol. 2017;13(3):183–93.

45. Geiger BC, Wang S, Padera RF, Grodzinsky AJ, Hammond PT. Cartilage-penetrating nanocarriers improve delivery and efficacy of growth factor treatment of osteoarthritis. Science Translational Medicine. 2018;10(469):eaat8800.

46. Zhou K, He X, Shi K, Hu D, Yang C, Gao M, et al. A dual responsive nanohydrogel system for sustained drug delivery and cartilage penetration in osteoarthritis therapy. Journal of Controlled Release. 2026;389:114494.

47. Jansen MP, Hodgins D, Mastbergen SC, Kloppenburg M, Blanco FJ, Haugen IK, et al. Can gait patterns be explained by joint structure in people with and without radiographic knee osteoarthritis? Data from the IMI-APPROACH cohort. Skeletal Radiology. 2024;53:2409 – 16.

48. Wang Y, Wu C, Tao J, Zhao D, Jiang X, Tian W. Differential proteomic analysis of tibial subchondral bone from male and female guinea pigs with spontaneous osteoarthritis. Exp Ther Med. 2021;21(6):633.

49. Allen KD, Mata BA, Gabr MA, Huebner JL, Adams SB, Jr., Kraus VB, et al. Kinematic and dynamic gait compensations resulting from knee instability in a rat model of osteoarthritis. Arthritis Res Ther. 2012;14(2):R78.

