## Supplementary figures and images for "sPLA_2_ inhibitor-loaded nanoparticles mitigate spontaneous osteoarthritis in guinea pigs"

### Supplementary Figure 1. Supplementary Video 1.Table 1.

#### Slide 1
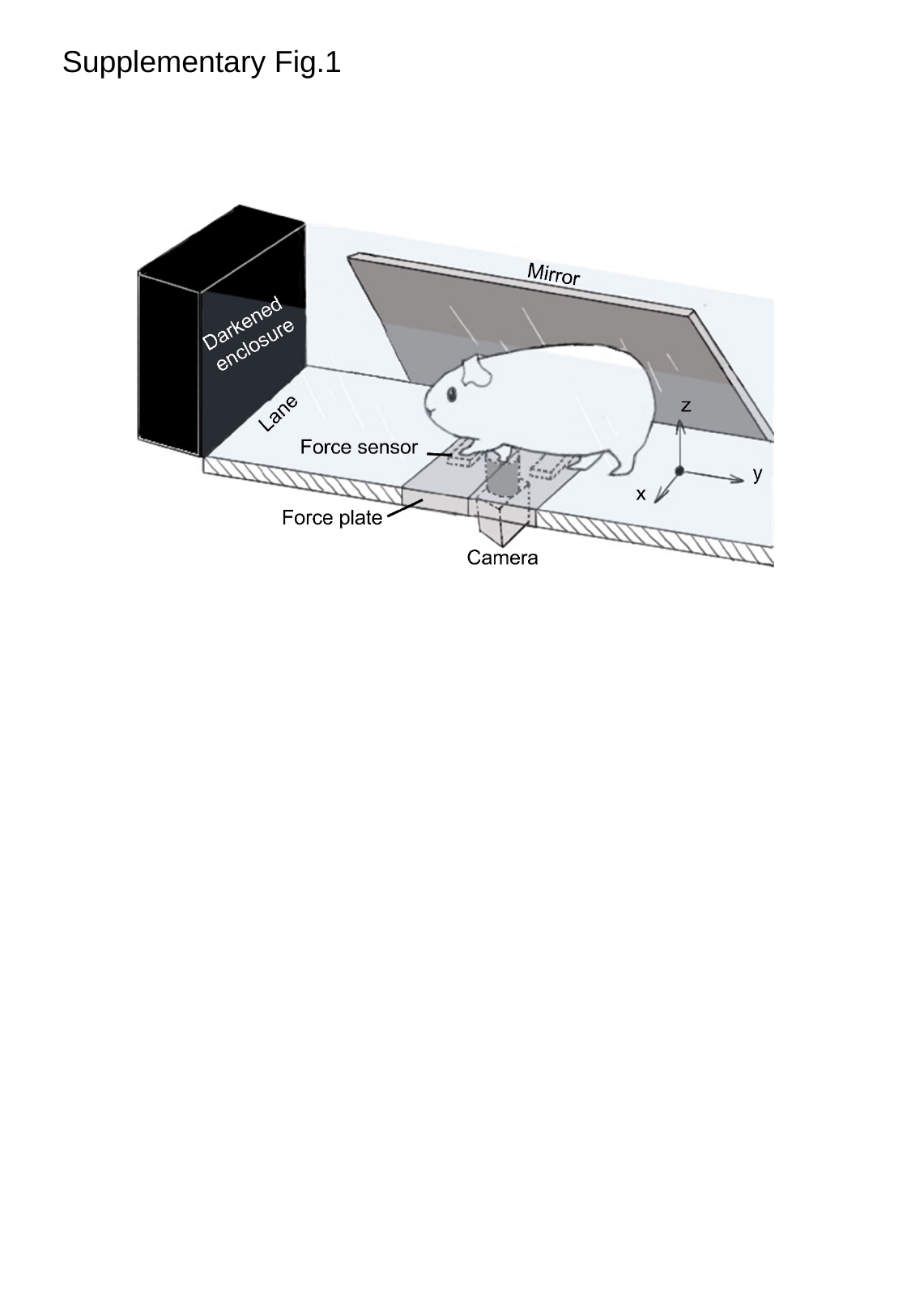

Supplementary Fig.1

#### Slide 2
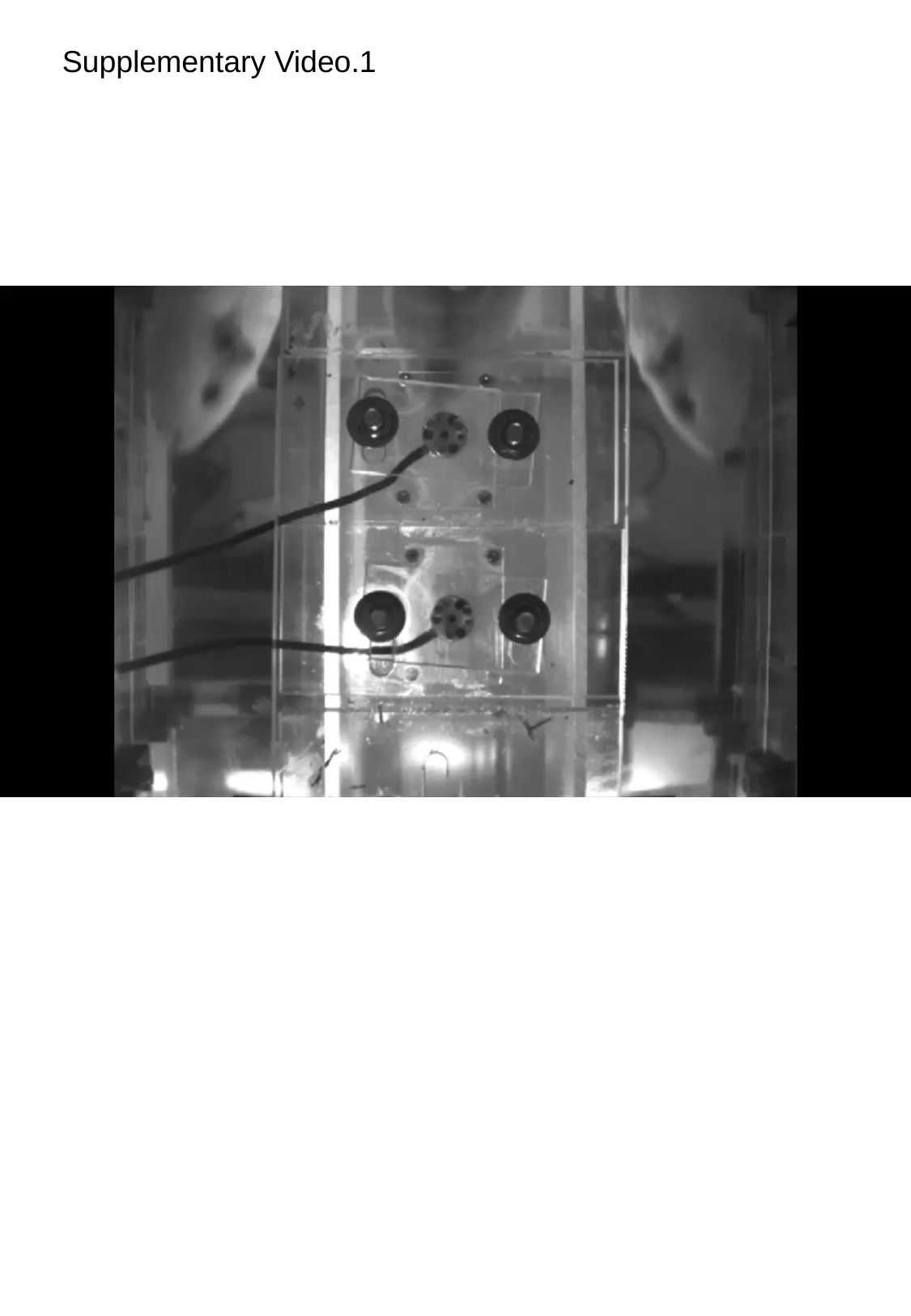

Supplementary Video.1

#### Slide 3
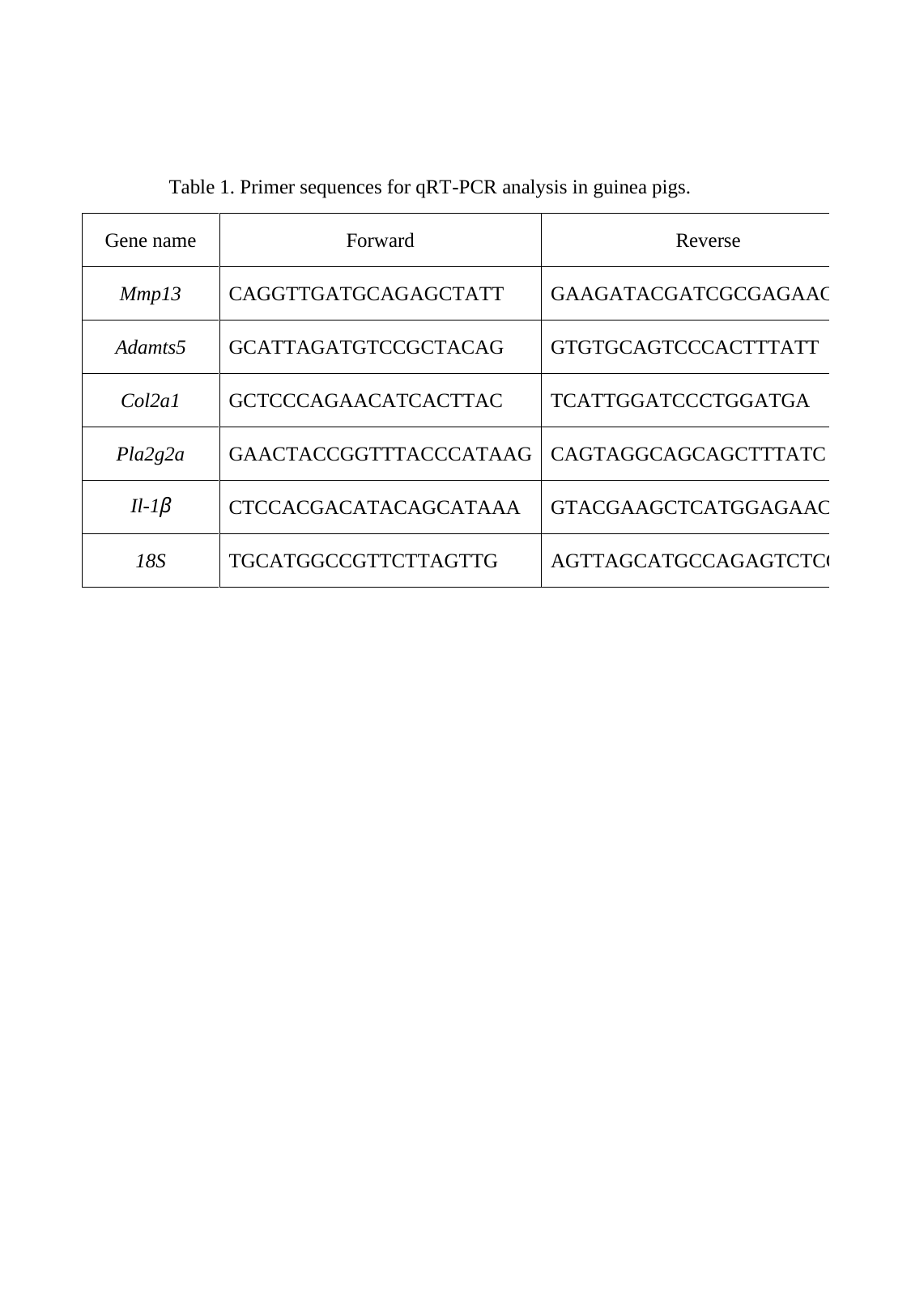
